# Global synthesis of aquatic insect heat tolerance reveals oxygen availability as a key driver of climate vulnerability

**DOI:** 10.64898/2026.03.23.713664

**Authors:** Stephanie Bristow, Wilco C.E.P. Verberk, Robby Stoks, Ben J. Kefford, Beatrice S. Dewenter, Alisha A. Shah

## Abstract

Accurately predicting species’ responses to climate change requires an understanding of the drivers of their thermal limits. Despite rapid warming of freshwater ecosystems worldwide, we still lack a global perspective on how upper thermal limits (UTLs) vary among aquatic insects, what constrains these limits, and how they contribute to species vulnerability. Here, we compiled a global dataset encompassing 423 aquatic insect species to test the effects of environmental conditions, organismal traits, acclimation history, and phylogenic relationships on patterns of heat tolerance. Maximum habitat temperatures were positively correlated with UTLs supporting the Climate Extremes Hypothesis, and insects relying exclusively on dissolved oxygen had the lowest UTLs supporting the Oxygen- and Capacity-Limited Thermal Tolerance hypothesis. Functional traits also explained substantial variation in UTLs; those that feed via scraping and shredding exhibited some of the lowest UTLs. Laboratory acclimation methods further influenced UTL estimates. Short-term exposure to higher acclimation temperatures increased UTLs, but longer exposure led to decreased heat tolerance. Finally, warming tolerance, i.e., the difference between UTL and the maximum habitat temperature) varied with breathing mode. Across latitude, warming tolerances were lowest for obligate dissolved oxygen-breathers but increased more rapidly in insects that can access terrestrial air. Collectively, these patterns indicate that oxygen is a key mechanism shaping thermal vulnerability in aquatic insects.

## Introduction

Ectotherm body temperatures reflect those of their environments (Angilletta, 2009), especially in aquatic ectotherms. Because all physiological functions are temperature-dependent, temperature has profound effects on survival, reproduction, and performance with consequences for populations and communities (Gilbert et al., 2014). The frequency of extreme heat events therefore threaten not only species persistence but also alter species abundances, distributions, and biotic interactions (Schleuning et al., 2020). Insects are declining at an alarming rate (Wagner, 2020) and evidence indicates that aquatic insects may be among the most severely impacted (Harvey et al., 2023). Despite their ecological importance and reported declines, we still know remarkably little about how sensitivity to heat stress varies among aquatic insects across broad geographic scales, and which environmental and organismal factors shape heat tolerance. Identifying the drivers of this variation can give us insight into the processes that shape sensitivity to heat in aquatic insects and for improving predictions of future responses to climate warming (Sunday et al., 2019).

Upper thermal limits (UTLs), which define the maximum temperature an organism can withstand prior to losing locomotor functionality (Bogert & Cowles, 1944; Lutterschmidt & Hutchison, 1997), are widely used metrics for measuring heat tolerance. As a result, UTLs have been key for linking physiology to environmental temperature and gaining insight into how variation in heat tolerance relates to latitudinal gradients which capture variation in temperature and seasonality (e.g., Deutsch et al. 2008; Sunday et al. 2019; Verberk et al., 2026). UTLs can be useful for broadly comparing species’ vulnerability to warming under climate change (Sunday et al., 2014). For example, one metric for vulnerability can be derived from UTLs by calculating the warming tolerance, i.e., the difference between mean UTL and the maximum habitat temperature a population experiences (Deutsch et al., 2008; Dewenter et al., 2025; Kingsolver et al., 2013). Moreover, proximity to upper thermal limits strongly predicts population-level responses to warming (Hamblin et al., 2017).

All species should evolve thermal limits that, at a minimum, match the highest and lowest temperatures typically experienced in their environment. The Climate Extremes Hypothesis (CEH) predicts that UTLs should be correlated with maximum environmental temperatures (Bozinovic, 2011; Pither, 2003). Thus, species exposed to higher maximum temperatures should evolve higher UTLs than those experiencing lower maximum temperatures (**Fig. 1A**). Macrophysiological studies of terrestrial and marine species largely meet these expectations: on land, species with the highest UTLs occur at mid-temperate latitudes and lower elevations; in the oceans, species with the highest UTLs occur in the tropics where annual maximum temperatures are highest (Deutsch et al., 2008; Polato et al., 2018; Sunday et al., 2010, 2019).

**Figure 1.**
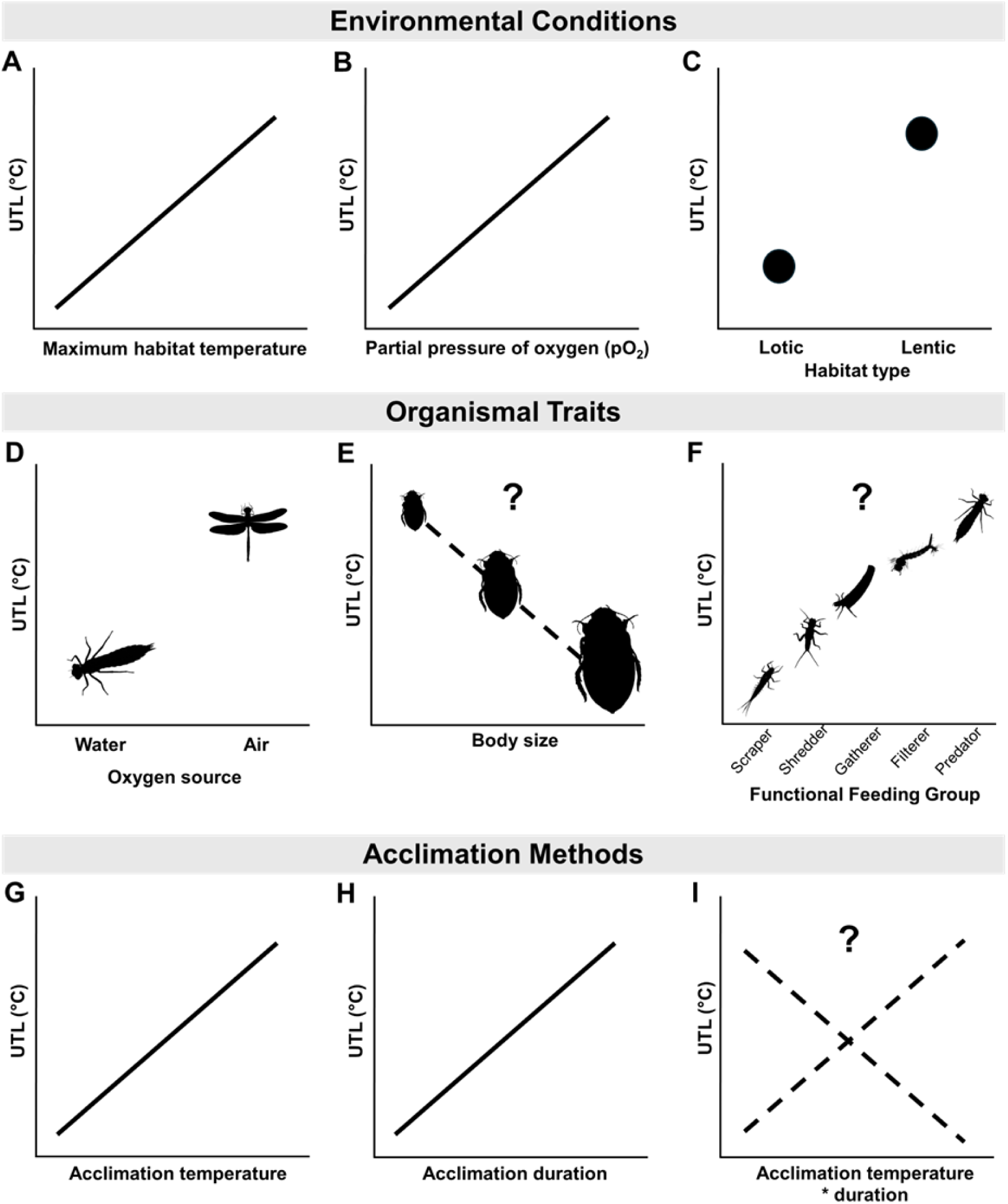
Predicted relationships between upper thermal limits (UTLs) and three non-mutually exclusive drivers: environmental conditions, organismal traits, and acclimation methods. **(A)** UTLs are expected to increase with higher maximum temperatures and **(B)** greater oxygen partial pressure. **(C)** UTLs may also be elevated in lentic habitats, which are typically warmer and often experience greater diel oxygen extremes relative to lotic systems. **(D)** Insects with access to atmospheric oxygen either through respiratory structure (i.e. breathing mode) or during terrestrial adult stages are predicted to exhibit higher UTLs. **(E)** The influence of body size and **(F)** functional feeding group on UTLs are unclear. Laboratory acclimation conditions are expected to shape UTL estimates, such that **(G)** higher temperatures and **(H)** longer acclimation durations will generally increase UTLs. However, the degree to which these factors interact **(I)** is unknown.

For aquatic insects, temperature is not the only driver of UTLs, and water oxygenation and flow are important modulators (Frakes et al., 2021; Verberk & Bilton, 2013). More broadly, core life processes and fitness depend on multiple abiotic factors, including temperature, oxygen, and water flow. However, water holds approximately 20-30 times less oxygen than air (Dejours, 1981; Verberk et al., 2011). This oxygen is difficult to extract due to the density and viscosity of water, making respiratory ventilation both challenging and energetically costly. Warming can exacerbate the challenge of extracting oxygen from the aquatic environment. This modulating effect of warming is encapsulated in the Oxygen- and Capacity-Limited Thermal Tolerance (OCLTT) hypothesis, which posits that rising temperatures can increase metabolic demand such that it outstrips oxygen supply imposing limits on insect tolerance to temperature (Pörtner, 2002; Pörtner & Knust, 2007; Verberk et al., 2011). One prediction that can be derived from the OCLTT hypothesis is that species with greater access to oxygen are predicted to tolerate higher temperatures (Verberk et al., 2016; **Fig. 1B & D**). Lentic systems (e.g., ponds and lakes) often experience warmer temperatures, thermal stratification, and episodic hypoxia, whereas lotic systems (e.g., streams and rivers) are more continuously mixed and aerated through flow (Frakes et al., 2021; Salem, 2021). Such habitat-level differences in thermal and oxygen dynamics may interact with organismal traits to shape heat tolerance (Verberk et al., 2018; **Fig. 1C**). Furthermore, aquatic insects have evolved respiratory adaptations to meet oxygen demands (Buchwalter et al., 2019; Skelly et al., 2002) and in well-oxygenated habitats, many rely on extracting dissolved oxygen via diffusion through the tegument or gills. In habitats with episodic low levels of dissolved oxygen, aquatic insects are adapted for aerial gas exchange. For example, mosquito larvae use breathing tubes (siphons) that directly access atmospheric air. Diving beetles periodically surface to renew the air supply carried under their elytra. Thus, the modulating effect of water oxygenation is likely different across habitat types and modes of breathing, explaining why some studies have found support for the OCLTT in aquatic insects (Frakes et al., 2021; Verberk et al., 2011), and others have not (Kim et al. 2017). Nevertheless, to what extent breathing mode and habitat type consistently impact UTLs across aquatic insects remains unresolved.

Higher oxygen demands in warming aquatic habitats may be especially challenging for large species (Leiva et al., 2019; Pauly, 2021; Verberk et al., 2011). For fish there is indeed evidence that larger species are more susceptible to oxygen limitation than smaller species, particularly in warm conditions (Navarro et al., 2025; Pörtner & Knust, 2007). However, despite well-established effects of body size on metabolic rate (Rubalcaba et al., 2020), and susceptibility to hypoxia (Deutsch et al., 2022; Verberk et al., 2022), the relationship between body size and heat tolerance seems small and variable (Dewenter et al., 2025; Leiva et al., 2019; Ospina & Mora, 2004). This suggests that species may have evolved many strategies to overcome such constraints. Thus, although it is likely that individuals of different sizes face distinct physiological challenges (Kraskura et al., 2023), how body size relates to upper thermal limits is not yet clear (**Fig 1E**).

Tolerance to heat stress may also be dependent on baseline metabolic demands associated with different feeding strategies. Aquatic insects are often grouped into functional feeding guilds (FFGs), which describe their function in the ecosystem based on feeding activity and the mechanisms used to consume food resources (Wallace & Webster, 1996). These include grazers (e.g., scrapers), shredders, gatherers, filterers, predators, and piercer-herbivores (Tomczyk et al., 2022). Minimum energy demands vary among guilds, especially when comparing guilds that require more movement (e.g., predators) versus those that are potentially more sedentary (e.g., filter feeders). In temperate mountain streams, for instance, predatory stoneflies had higher resting metabolic rates compared to grazing mayflies from the same streams (Shah et al., 2021). Thus, one possibility is that guilds with higher metabolic demands may incur greater costs under warming than those with lower demands (Boyero et al., 2012). Alternatively, species with higher metabolic demands may also display higher mobility and ability to thermoregulate (**Fig. 1F**) (Poff et al., 2006; Tomczyk et al., 2022). To our knowledge, no studies have investigated the joint effects of feeding guilds and other abiotic and biotic variables on global patterns of aquatic insect UTLs (Hering et al., 2009; Pyne & Poff, 2017; Tomczyk et al., 2022), leaving an important gap in our understanding of which groups overall may be more sensitive to warming.

Interpreting the environmental and biological drivers of UTLs is further complicated by the lack of methodological standardization across studies, which undermines direct comparisons of reported values (Santos et al., 2011; Terblanche et al., 2007). A commonly used method of estimating UTLs is measuring the critical thermal maximum (CT_MAX_) in which temperature is ramped at a consistent rate until the animal loses the ability to right itself or is otherwise unable to control its movements (Lutterschmidt & Hutchison, 1997). Animals in this state are unable to function in their environment and are generally considered ecologically dead (Kingsolver & Umbanhowar, 2018; Lutterschmidt & Hutchison, 1997). In aquatic insects, CT_MAX_ is indicated by a loss of righting (e.g., Dallas and Rivers-Moore) often preceded by muscular spasms or movement to the water’s surface (Shah et al. 2017). A second approach to estimating UTLs is by measuring lethal temperatures (LTs), which exposes animals to a static temperature for a fixed duration to determine the temperature at which there is 50 % (LT_50_) or 100 % (LT_100_) mortality (Huey & Stevenson, 1979; Lutterschmidt & Hutchison, 1997). Whereas CT_MAX_ is a metric of response to acute heat stress, LTs can shed light on responses to prolonged, chronic warming. Although recent global syntheses of UTLs span an array of taxa (Bennett et al., 2021; Sunday et al., 2010, 2019) they often exclude LTs (Chown et al., 2015; Weaving et al., 2022) likely because static and dynamic temperature assays are not directly comparable. However CT_MAX_ and LTs are mathematically related (Arnold et al., 2025; Ørsted et al., 2022; Rezende et al., 2014), and using statistical correction to account for methodological differences among assay types, can expand taxonomic coverage (Molina et al., 2025; Verberk et al., 2026).

Finally, UTLs are not necessarily fixed – many species exhibit physiological plasticity in thermal limits with laboratory acclimation prior to experiments (Gunderson & Stillman, 2015; Havird et al., 2020; Weaving et al., 2022). Although excellent recent reviews have shown that UTLs are generally less plastic than lower thermal limits (Gunderson & Stillman, 2015; Weaving et al., 2022), incorporating laboratory conditions such as acclimation temperature and duration can allow us to better interpret and compare UTL values across different studies. Acclimation temperature and duration of exposure may interact (**Fig. 1G, H & I)** and affect whether organisms display short-term heat hardening or have decreased UTLs because of accumulated heat injury (Weaving et al., 2022; Wehrli et al., 2024). It is, therefore, essential to address and distinguish methodological effects from environmental and organismal drivers of heat tolerance in large comparative analyses.

Here, we synthesize a global dataset using a hierarchical analytical framework that accounts for phylogenetic non-independence and methodological heterogeneity to test *a priori* hypotheses regarding the independent and interactive effects of environmental conditions, organismal traits, and acclimation history on UTLs in aquatic insects. We explicitly model abiotic and biotic drivers jointly rather than in isolation, enabling identification of shared underlying mechanisms and reducing the risk of missing influential predictors that emerge only through their covariation. Specifically, we evaluate how thermal and hypoxic conditions interact with key organismal traits — including breathing mode, life stage, and habitat type — that govern oxygen acquisition and heat stress tolerance. By integrating these drivers across a phylogenetically diverse dataset, we evaluate competing theoretical frameworks, including the CEH and the OCLTT hypothesis, linking mechanistic predictions to empirical variation in UTLs and generating quantitative estimates of vulnerability in aquatic insects.

## Materials and Methods

### Literature Search

PRISMA guidelines were used for reporting each step of the literature review process (Page et al., 2021). We included the following Google Scholar search terms: ((ALL=(aquatic insect OR stream insect)) AND ALL=(thermal tolerance OR thermal limit OR critical thermal maximum OR CT_MAX_ OR lethal)) We included the following Web of Science search terms: ((aquatic insect OR stream insect) AND ALL=(thermal tolerance OR thermal limit OR critical thermal maximum OR CT_MAX_ OR lethal)). These queries resulted in 82,100 and 1,649 results in Google Scholar and Web of Science, respectively. Most sources from earlier datasets including aquatic insect UTLs were also captured in these search results (Bayat et al., 2025; Bennett et al., 2021; Chown et al., 2015; Gunderson & Stillman, 2015; Leiva et al., 2019; Seebacher et al., 2015; Weaving et al., 2022). The search procedure for eligible sources in our analysis is also reported in **Fig. S1** (**Appendix 1 in Supporting Information**). After assembling all journal articles from existing datasets, we screened for additional eligible sources by reviewing abstracts. Studies were included in our analysis only if they measured aquatic insect upper thermal limits (CT_MAX_, and upper LT_50_ or LT_100_) using either static or dynamic ramping methods. Subsets of data from eligible sources were excluded if they included other sources of variation (e.g., hypoxia, salinity, or toxins). We also included four sources of unpublished UTL data from research conducted by authors of this manuscript.

### Data extraction

We recorded thermal limit measurements and error estimates from source tables or text when reported. If results only appeared in figures, mean and error estimates were measured from plots using ImageJ (Schneider et al., 2012). We also converted median values to mean estimates using sample sizes and quartiles or individual-level data when available.

### Moderator variables

#### Environment: Geographic features & climate

We included a number of additional variables for each location where insects in our dataset were found, including latitude, longitude, elevation, habitat type, maximum habitat temperatures, and pO_2_. Habitat types were assigned based on site descriptions, where streams, creeks, and rivers were categorized as “lotic,” and all still waters including lakes, and ponds were categorized as “lentic”. If studies reported latitude, longitude and elevation, these were recorded; otherwise, we approximated these parameters based on site or larger locality names, such as towns, cities, or provinces.

Elevation primarily reflected collection locations: however, if organisms were collected at higher elevations and subsequently tested elsewhere, we used estimated elevation of the test location when available. Because elevations of testing sites were not consistently reported across studies, elevation estimates represent a mixture of collection and test conditions. Where both were available, values were generally similar (median difference = 0 m; r = 0.73), indicating that the combined elevation variable provides a reasonable approximation of elevation conditions at collection sites, including thermal regime and atmospheric pressure.

Using coordinates of collection sites, we extracted maximum habitat temperatures in two ways for aquatic insects tested in water or in air, i.e., during terrestrial life stages. For organisms tested in water, we extracted surface temperatures of aquatic environments with the FutureStreams modeled water-temperature climatology, using the averaged maximum water temperature of the hottest month from the E2O (1979–2005) historical forcing period (Bosmans et al., 2022). For insects tested in air, we extracted averaged maximum air temperature of the hottest month from WorldClim v2.1, via BIO5, and the 1970–2000 baseline (Fick & Hijmans, 2017). We also included partial pressure of oxygen (pO_2_) by first estimating atmospheric pressure at the elevation (meters above sea level) for each site using the standard barometric formula (P = 101.325 × (1 – 2.25577 × 10⁻⁵ × elevation)⁵·²⁵⁵⁸, in kPa) and then calculating pO_2_ as 20.95% of atmospheric pressure (P).

#### Organismal Traits

For each unique species in our dataset, we recorded breathing mode, i.e., the way an insect extracts oxygen from the environment. These modes included diffusion through cuticles and gills, the use of plastrons and siphons, as well as bimodal breathing (i.e., the use of both gills and atmospheric air through spiracles), and finally, spiracles. We also recorded organisms’ primary functional feeding groups, including scrapers, shredders, gatherers, filterers, predators, and piercer-herbivores, based on information from Kefford et al. (2020) and Merritt and Cummins (1995). Surprisingly, very few studies measured and reported body sizes; therefore, body size analyses were performed on a subset of data including wet or dry mass (mg) measurements. To reduce the broad range of body sizes, all mass values were transformed on a log_10_ scale.

#### Experimental methods

Variables recorded for each study’s experimental methods included the following: type of thermal assay (static or dynamic ramping), type of thermal limit (i.e., CT_MAX_, LT_50_, LT_100_), starting temperature, ramping rate (°C/minute), acclimation temperature, and acclimation duration. For dynamic assays, we calculated time to the thermal endpoint (minutes) as (final - starting temperature) divided by ramping rate. Acclimation temperature covaried with environmental temperature in our dataset; therefore, we used standardized acclimation temperatures (Δ°C; acclimation temperature - species-specific mean acclimation temperature) to isolate within-species effects of acclimation temperature on thermal limits. Acclimation periods ranged from hours to weeks across studies, so we transformed acclimation duration (hours) to a log_10_ scale to reduce the scale for analysis. <u>Statistical Analyses</u>

#### UTL correction

Insects tested in dynamic versus static assays were exposed to different durations and intensities of thermal stress, producing differences in UTL estimates that were not directly comparable. We therefore applied a time-standardized statistical correction to thermal limits. First, we regressed each thermal limit against time to the thermal endpoint on a log_10_ scale. We used this first model to predict thermal limits and used a second model to estimate standardized limits corresponding to a 100-minute exposure. The difference between the first model’s predicted thermal limits and the second model’s standardized limits was applied to each raw thermal limit to obtain corrected UTL values (refer to **Appendix 2** for model results).

Before correction, UTLs from static assays were 8.6 °C lower than those from dynamic assays (F₁,₁₃₄₄ = 350.2, p < 0.001; R² = 0.21). After applying the duration-based correction, the difference between assay types was reduced to 1.37 °C, a sixfold decrease, and assay type explained <1% of the variation in UTLs (F₁,₁₃₄₄ = 9.30, p = 0.002, R²=0.01). Thus, the correction substantially reduced methodological bias, although a small but significant difference remained (refer to **Appendix 2** for statistical comparisons before and after correction). Assay types were unevenly distributed across insect orders and primary oxygen sources (refer to **Appendix 3** for reported distributions of assay types). Corrected UTL values were used as the response variable in subsequent analyses. Across all species, UTLs ranged from 16.3 to 55.8 °C with a median of 34.2 °C and mean of 35.5 °C.

#### Phylogeny

To account for non-independence among species due to shared evolutionary history, we incorporated phylogenetic relationships as a random effect in phylogenetic generalized linear mixed models (PGLMMs), following the approach described in Dewenter et al. 2025. Briefly, species were matched to the Open Tree of Life using the R packages *rotl*, *caper*, and *ape* (Michonneau et al., 2016; Orme, 2023; Paradis et al., 2004), and an induced subtree was generated to estimate branch lengths. For taxa not represented in Open Tree of Life or fully identified to species in the original article, a placeholder was added for potential unique species occupying different regions (e.g., “sp. 1”) and added as a descendant of the corresponding genus node. Phylogenetic signal in UTLs was independently assessed using Pagel’s λ (ranging from 0-1) with *phytools* (Revell, 2012).

#### PGLMMs

Because 18.9 % of species appeared in more than one article, and 56 % had more than one UTL estimate, we used phylogenetic generalized linear mixed models (PGLMMs) from the R package *phyr* (Li et al., 2020) to incorporate intraspecific variation in thermal limits. We compared each model’s performance using their likelihood, AIC and BIC, and R^2^. We first structured GLMs without phylogenetic correction, including both an intercept-only and insect Order model, before implementing an intercept only PGLMM. Then, to evaluate how UTLs vary with geographic and environmental factors, we compared two sets of PGLMMs either including absolute latitude, hemisphere, and elevation or maximum habitat temperature, and partial pressure of oxygen (pO_2_). We compared these models to determine whether environmental proxies (latitude & elevation) can match performance of direct environmental variables (habitat temperature & pO_2_). Next, including the abiotic moderators of the best performing model of this set, we ran three new models adding biotic variables: breathing mode, functional feeding group, and both traits. Finally, we extracted residuals from the best performing PGLMM including abiotic and biotic predictors to compare UTLs between life stages, habitat types, body mass, and acclimation variables (standardized acclimation temperature, acclimation duration, and their interaction). For models comparing UTLs across aquatic habitat types, data from terrestrial adult stages were omitted, and included adults when they were fully aquatic. To complement these analyses with phylogenetically corrected residuals, we also fit general linear models (LMs) using UTLs for each predictor set (life stage, habitat type, body mass, and acclimation variables). These models were used primarily to assess whether detected relationships in the phylogenetically corrected analyses were also apparent without phylogenetic correction or additional abiotic and biotic moderators (refer to **Appendices 6-9** for life stage, habitat type, body mass, and acclimation variables GLM results).

#### Warming tolerance

Following Kingsolver, Diamond, and Buckley (2013) and Sunday et al. (2014) we calculated warming tolerance as: UTL – maximum habitat temperature to evaluate how close species are living to their UTL based on maximum temperatures of their environments. We structured a separate linear model (**Appendix 10**) and phylogenetically corrected model to estimate how warming tolerance changed with absolute latitude and interacted with primary source of oxygen (i.e., DO in water, or air).

## Results

Our search revealed 96 unique and original sources meeting our criteria for measuring aquatic insect upper thermal limits using standard methods and un-pooled datasets. Our dataset included 8 Orders, 232 genera, and 423 species of aquatic insects (**Fig. 2A & B**). Large parts of the globe were not represented in our dataset because, to our knowledge, no studies fitting our criteria have been conducted there. These areas included parts of the American tropics, and much of Asia, Africa, and South America. We analyzed a total of 1,346 mean estimates of UTLs across dynamic ramping (n=1,127), and static (n = 219) assays.

**Figure 2.**
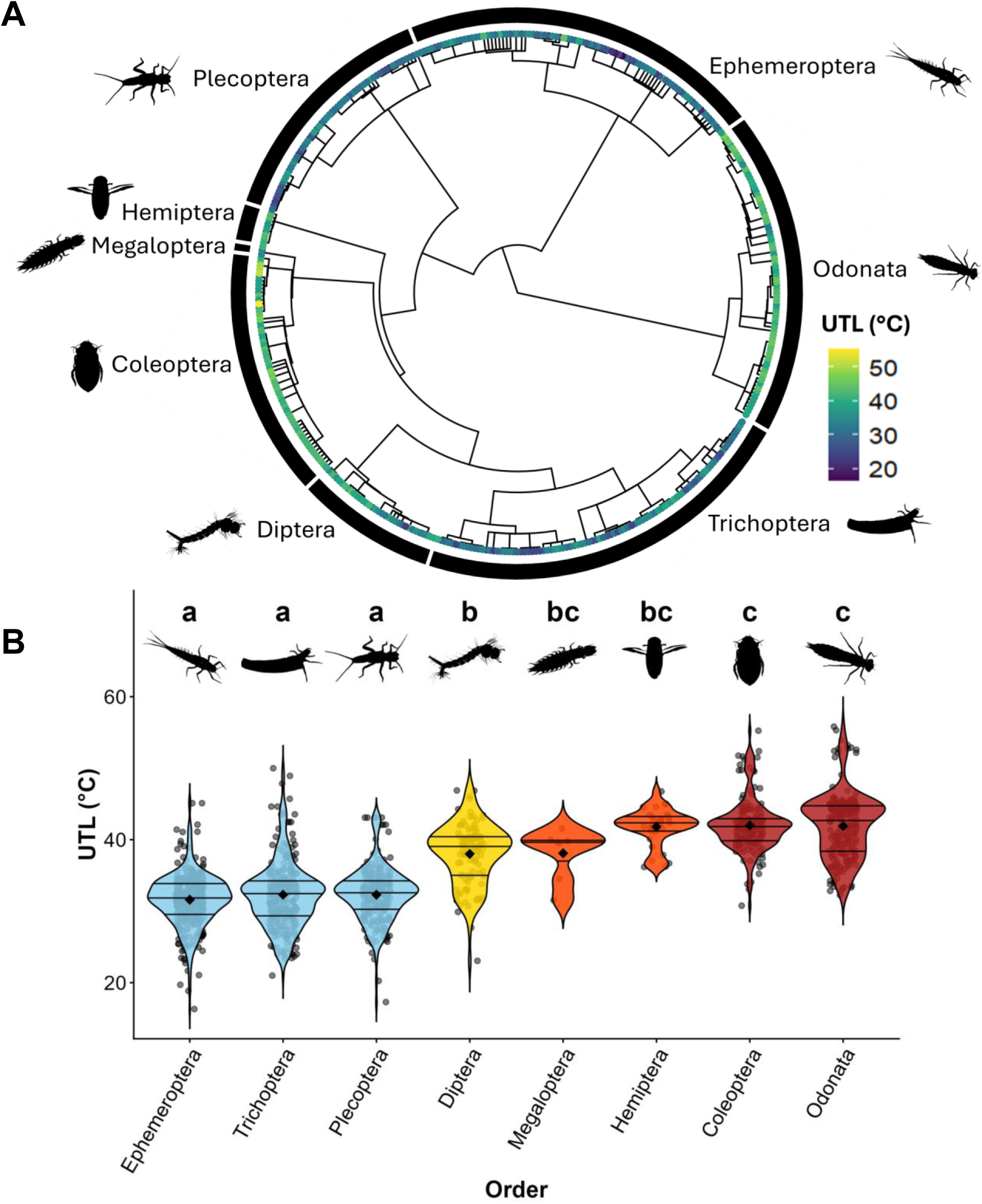
(A) Upper thermal limits (UTL) are presented across the phylogenetic tree for all species in our dataset and lineages are outlined within their respective taxonomic order. **(B)** UTLs are grouped by order and include observed UTLs (points) with model derived means (diamonds) and quantiles, with statistically similar groups presented with the same color and letters.

### Model comparisons

UTLs displayed strong phylogenetic signal with a high Pagel’s λ (λ = 0.89, p < 2x10⁻¹⁶). Thus, accounting for phylogenetic relatedness significantly improved model performance (**Table 1**). The non-phylogenetic model with taxonomic order explained moderate variation in UTLs (R² = 0.55) but performed worse than a phylogenetically informed intercept-only PGLMM (R²𝖫ᵢₖ = 0.72; R²_pred_ = 0.75). This improvement supported the use of PGLMMs for subsequent analyses.

**Table 1.**
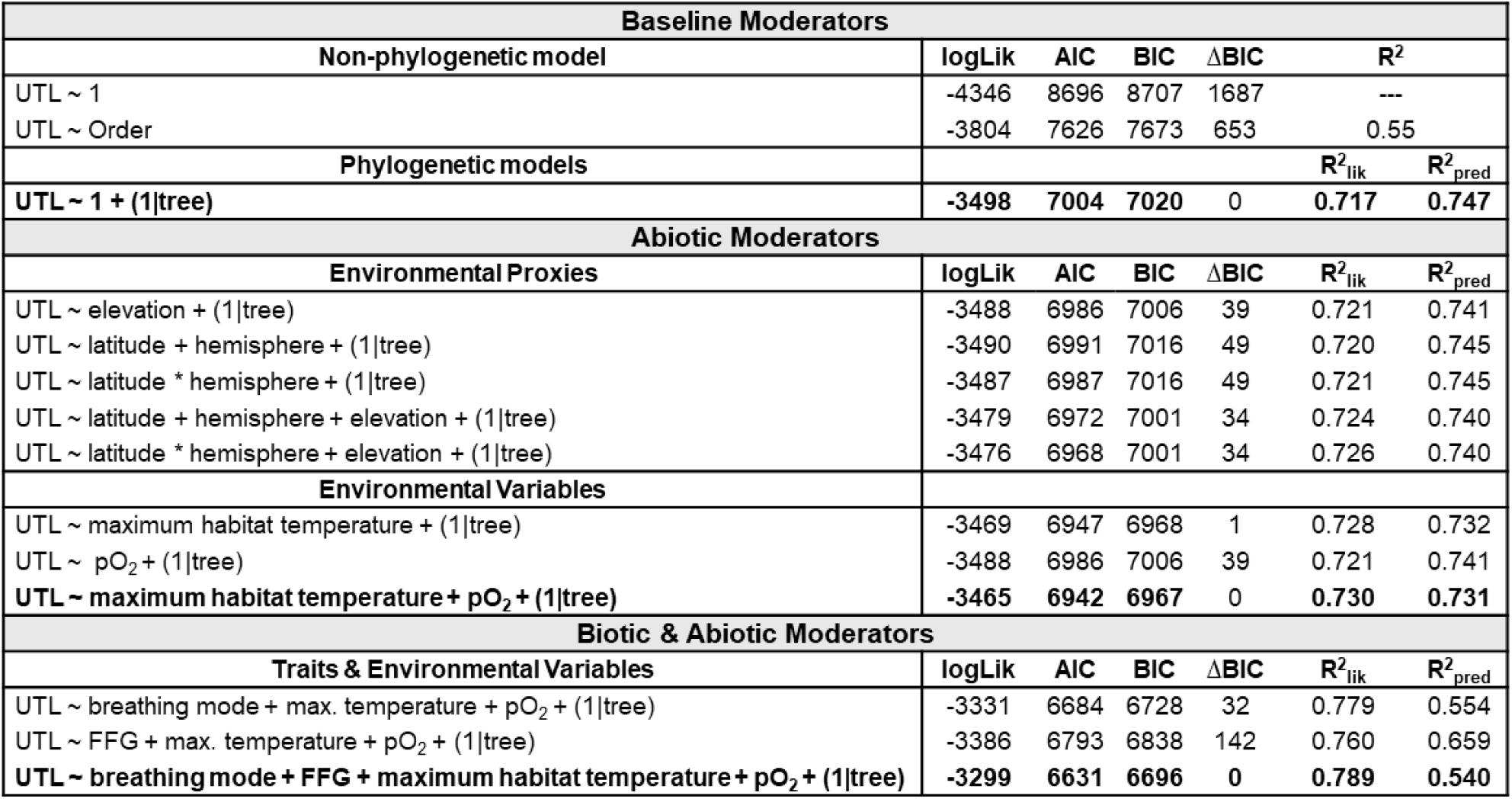
Comparisons of general linear and phylogenetic generalized linear mixed models (PGLMMs). Log likelihood, AIC, BIC, ΔBIC, and predictive performance metrics **R^2^lik** and **R^2^pred** are used to compare model performance

For abiotic explanatory variables, models including elevation, absolute latitude, and hemisphere had similar explanatory power, but the model including maximum habitat temperature and partial pressure of oxygen (pO₂) received stronger support (**Table 1**). Adding biotic moderators further improved model fit, and the model with maximum habitat temperature, pO₂, breathing mode, and FFG was the best supported model with abiotic and biotic moderators (**Table 1**). However, its predictive performance (R²_pred_ = 0.54) was lower than that of models with only abiotic moderators (R²_pred_ = 0.73). This reduction likely reflects how breathing modes and functional feeding groups are to some extent phylogenetically structured and additionally covary with abiotic variables, introducing overlapping information that reduces the model’s ability to generalize patterns across species.

### Environmental Variables

Maximum habitat temperature and pO_2_ predictively explained variation in UTLs (**Table 2**, **Fig. 3A & B**). According to the model including maximum habitat temperature and pO_2_, a 1 °C increase in maximum habitat temperature corresponded to a 0.18 ± 0.03 °C increase in UTLs (Z= 6.41, p < 0.001). After accounting for maximum habitat temperature, pO_2_ unsurprisingly accounted for little additional variation in UTLs, as dissolved oxygen is correlated with temperature (Z = 1.46, p = 0.145). After including breathing mode and FFG, maximum habitat temperature remained a significant predictor (0.09 ± 0.03, Z = 3.04, p = 0.002).

**Figure 3.**
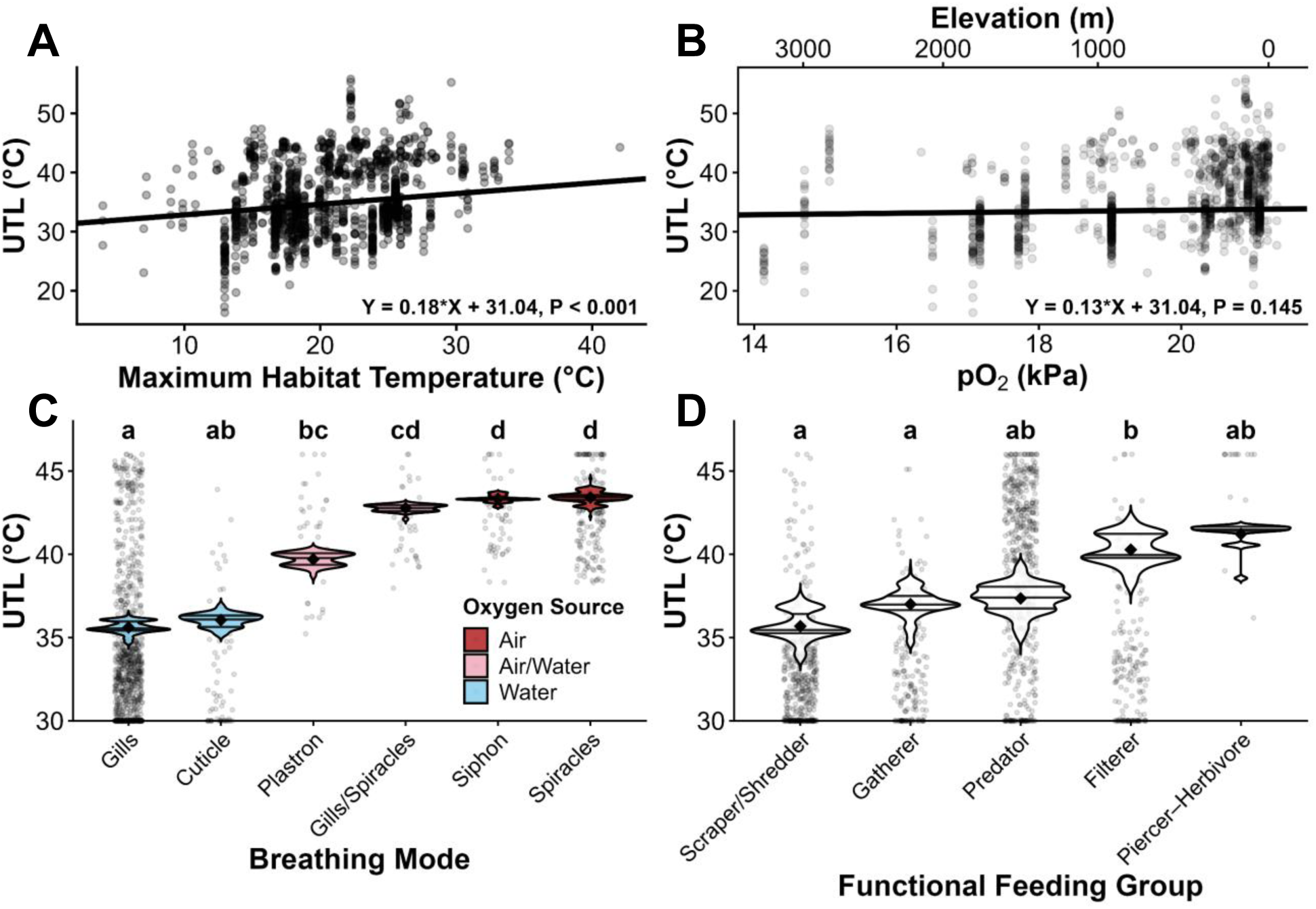
Observed UTLs (points) and phylogenetic generalized linear mixed model-fitted relationships (lines) between UTLs and **(A)** maximum habitat temperature and **(B)** oxygen partial pressure (pO₂). A secondary x-axis indicates the elevation (m) used to derive pO₂. Panels **C** and **D** show observed UTLs and overlain violin plots of PGLMM-predicted UTLs by trait group; violins and diamonds represent group-level predicted distributions and means, respectively (predictions retain the underlying ranges of maximum temperature and pO₂ experienced by each group). Letters denote significant differences in UTLs across **(C)** breathing modes (shaded by oxygen source) and **(D)** functional feeding groups.

**Table 2.** Results from LMs fitted to phylogenetically corrected residuals from intercept-only and full PGLMMs including environmental and trait moderators, examining effects of life stage, body size, habitat type, and acclimation variables

| Predictor | Residual Model | Variable | Estimate | S.E. | t | p-value |
| --- | --- | --- | --- | --- | --- | --- |
| Life stage | Intercept only<br>PGLMM | Juvenile (Intercept) | -0.19 | 0.076 | -2.451 | 0.014 |
|  |  | Adult | 1.18 | 0.182 | 6.476 | <0.001 |
|  | Full PGLMM | Juvenile (Intercept) | 0.06 | 0.069 | 0.835 | 0.404 |
|  |  | Adult | -0.12 | 0.167 | -0.746 | 0.456 |
| Body size | Intercept only<br>PGLMM | Intercept | -0.65 | 0.323 | -1.964 | 0.051 |
|  |  | Log Dry Mass (mg) | 0.27 | 0.138 | 1.963 | 0.051 |
|  |  | Log Wet Mass (mg) | -0.16 | 0.322 | -1.333 | 0.184 |
|  | Full PGLMM | Intercept | -0.69 | 0.310 | -2.216 | 0.028 |
|  |  | Log Dry Mass (mg) | 0.21 | 0.129 | 1.625 | 0.106 |
|  |  | Log Wet Mass (mg) | 0.18 | 0.302 | -0.111 | 0.911 |
| Habitat type | Intercept only<br>PGLMM | Lotic (Intercept) | -0.06 | 0.078 | -0.807 | 0.420 |
|  |  | Lentic | 0.03 | 0.176 | 0.176 | 0.860 |
|  | Full PGLMM | Lotic (Intercept) | 0.01 | 0.071 | 0.154 | 0.878 |
|  |  | Lentic | 0.32 | 0.160 | 1.996 | 0.046 |
| Acclimation | Intercept only<br>PGLMM | Intercept | 0.80 | 0.251 | 3.198 | 0.001 |
|  |  | Acclimation Treatment | 0.29 | 0.067 | 4.326 | <0.001 |
|  |  | Acclimation Duration | -0.47 | 0.141 | -3.334 | <0.001 |
|  |  | Acclimation Treatment * Duration | -0.11 | 0.038 | -2.778 | 0.006 |
|  | Full PGLMM | Intercept | 0.66 | 0.241 | 2.754 | 0.006 |
|  |  | Acclimation Treatment | 0.24 | 0.064 | 3.685 | <0.001 |
|  |  | Acclimation Duration | -0.33 | 0.135 | -2.414 | 0.016 |
|  |  | Acclimation Treatment * Duration | -0.08 | 0.036 | -2.292 | 0.022 |

### Organismal Traits

#### (a) Breathing mode

Breathing mode strongly predicted UTLs (**Fig. 3C**). Insects with access to atmospheric oxygen (siphon- & spiracle-breathers) exhibited significantly higher UTLs than species dependent on dissolved oxygen (insects breathing through gills and cuticles). When breathing mode was included in the PGLMM, the effect of pO₂ became statistically weaker, indicating that differences in oxygen acquisition mediates the relationship between oxygen availability and UTLs (refer to **Appendix 5** for breathing mode comparisons from PGLMM estimates and partial residuals).

#### (b) Functional Feeding Group (FFG)

Including functional feeding group (FFG) notably increased model performance (Table 2, **Fig. 3D**), yet many groups overlapped in significance (refer to **Appendix 5** for FFG comparisons from PGLMM estimates and partial residuals). Piercer-Herbivores exhibited some of the highest UTLs. Scrapers/shredders exhibited some of the lowest UTLs and were statistically distinct from filterers. However, scrapers/shredders, predators, and gatherers, displayed consistent overlap in their UTLs.

### Residual models

#### (a) Life stage

Results from a general linear model showed that adults had 8.65 ± 0.37°C higher UTLs than juveniles (**Appendix 6, Table S6:** p < 0.001). This pattern was also supported by the LM fit on intercept-only PGLMM residuals but was lost after accounting for abiotic and biotic moderators with the full PGLMM’s residuals (**Table 2**).

#### (b) Body size

Body size showed inconsistent associations with UTLs across analyses. LMs performed on UTL values detected positive relationships for both wet (**Appendix 7**, **Table S7**: p = 0.03) and dry mass (**Table S7**: p < 0.001) but likely represent interspecific differences in size. To evaluate within-species effects, we fit a linear mixed-effect model with species identity as a random intercept. In this model, wet mass no longer predicted UTLs (**Table S8**: p = 0.272), but dry mass remained statistically significant (**Table S8**: p < 0.001). These results were still based on a smaller dataset (52 species) and could still reflect interspecific variation rather than a consistent within-species mechanism.

Neither wet nor dry mass explained variation in PGLMM residuals (**Table 2**). After accounting for phylogenetic relationships and abiotic and biotic moderators, body size contributed little to no explanatory power. Overall, our analyses indicate that body-size effects on UTLs are weak and may be limited by low within-species sampling.

#### (c) Habitat type

LM results revealed that insects from lentic habitats had 8.08 ± 0.35°C higher UTLs than lotic species on average (**Appendix 8, TableS9:** p < 0.001). However, the statistical significance was largely reduced in the LMs fit on PGLMM residuals (**Table 2**).

#### (d) Acclimation

Standardized acclimation temperature and acclimation duration consistently affected UTLs according to LMs fit on non-phylogenetically corrected data (**Table 2, Appendix 9**) and PGLMM residuals. Acclimation at temperatures above a species’ mean acclimation temperature showed a positive effect on UTLs, while acclimation duration (log_10_ hours), and the interactive effects of standardized acclimation temperature and duration both negatively influenced UTLs (**Table 2**, **Fig. 4**). Generally, UTLs were highest with short-term exposure to higher acclimation temperatures (indicating potential for beneficial acclimation with brief heat shocks), but longer acclimation durations led to lower UTLs for species acclimated above or below their species’ mean acclimation temperatures.

**Figure 4.**
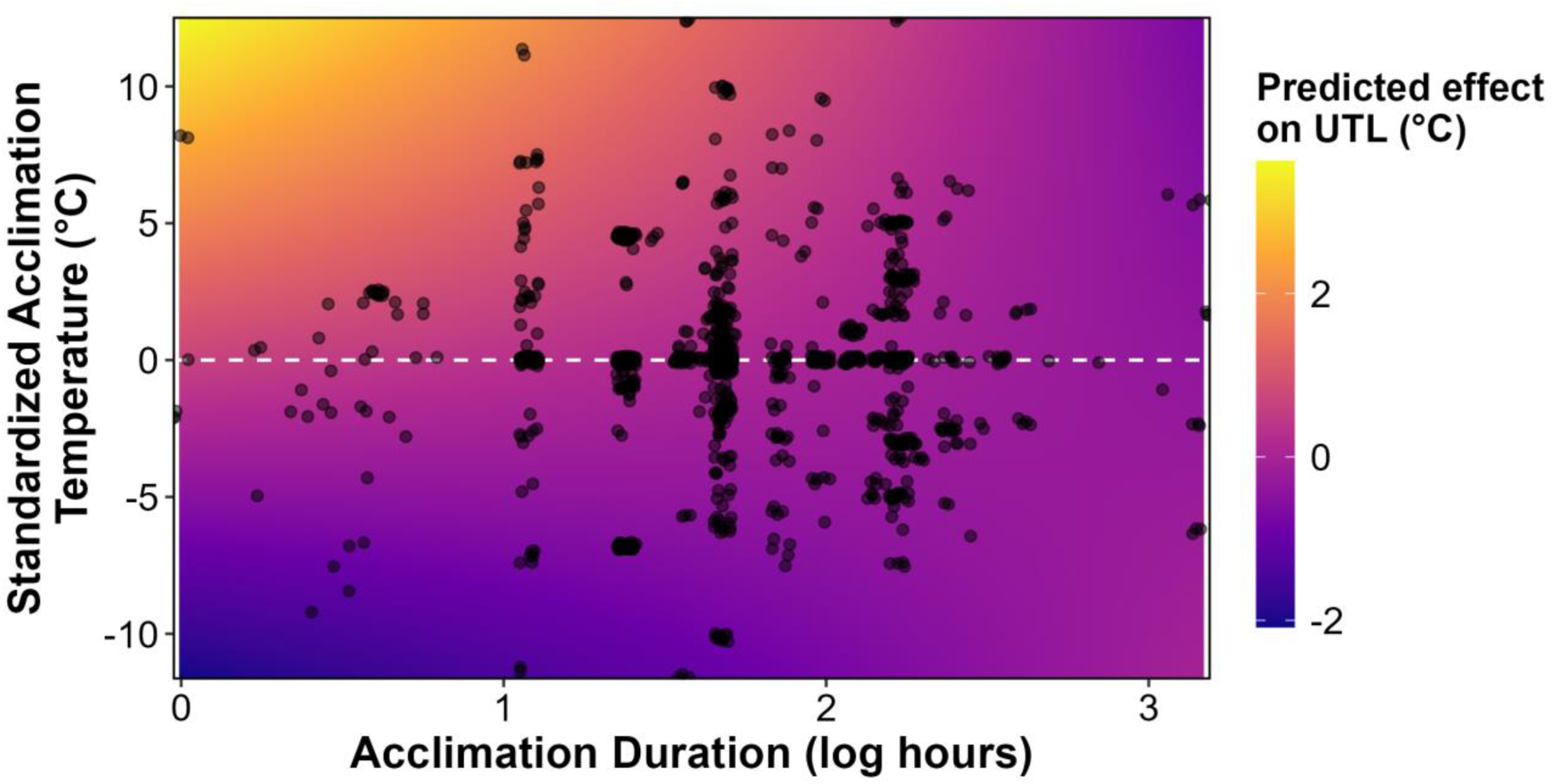
Predicted effects of acclimation variables on upper thermal limits (UTL,°C), estimated from residual variation after accounting for maximum habitat temperature, pO_2_, breathing mode, & functional feeding group. Standardized acclimation temperature (°C) was calculated as acclimation temperature - the species-specific mean temperature; the dashed horizontal line indicates no deviation from the species’ mean. Acclimation duration is shown on a log_10_ scale (hours). Points represent observations, and the background color scale indicates the predicted effect of acclimation on UTLs.

#### Warming tolerance

At the equator, warming tolerance (°C; UTL – maximum habitat temperature) did not differ significantly between dissolved oxygen or air breathers (p = 0.374). Warming tolerance increased significantly with absolute latitude, but the strength of this relationship differed between air and dissolved oxygen breathing insects (**Table 3**, **Fig. 5B**). Dissolved-oxygen breathing insects gain 0.15 ± 0.02 °C of warming tolerance per 1° increase in absolute latitude (p < 0.001), while air-breathers gain 0.21 ± 0.03 °C warming tolerance per 1° increase in absolute latitude (p = 0.065). These patterns of warming tolerance by oxygen sources were strongly supported by the LM fit to non-phylogenetically corrected data (**Appendix 10**).

**Figure 5.**
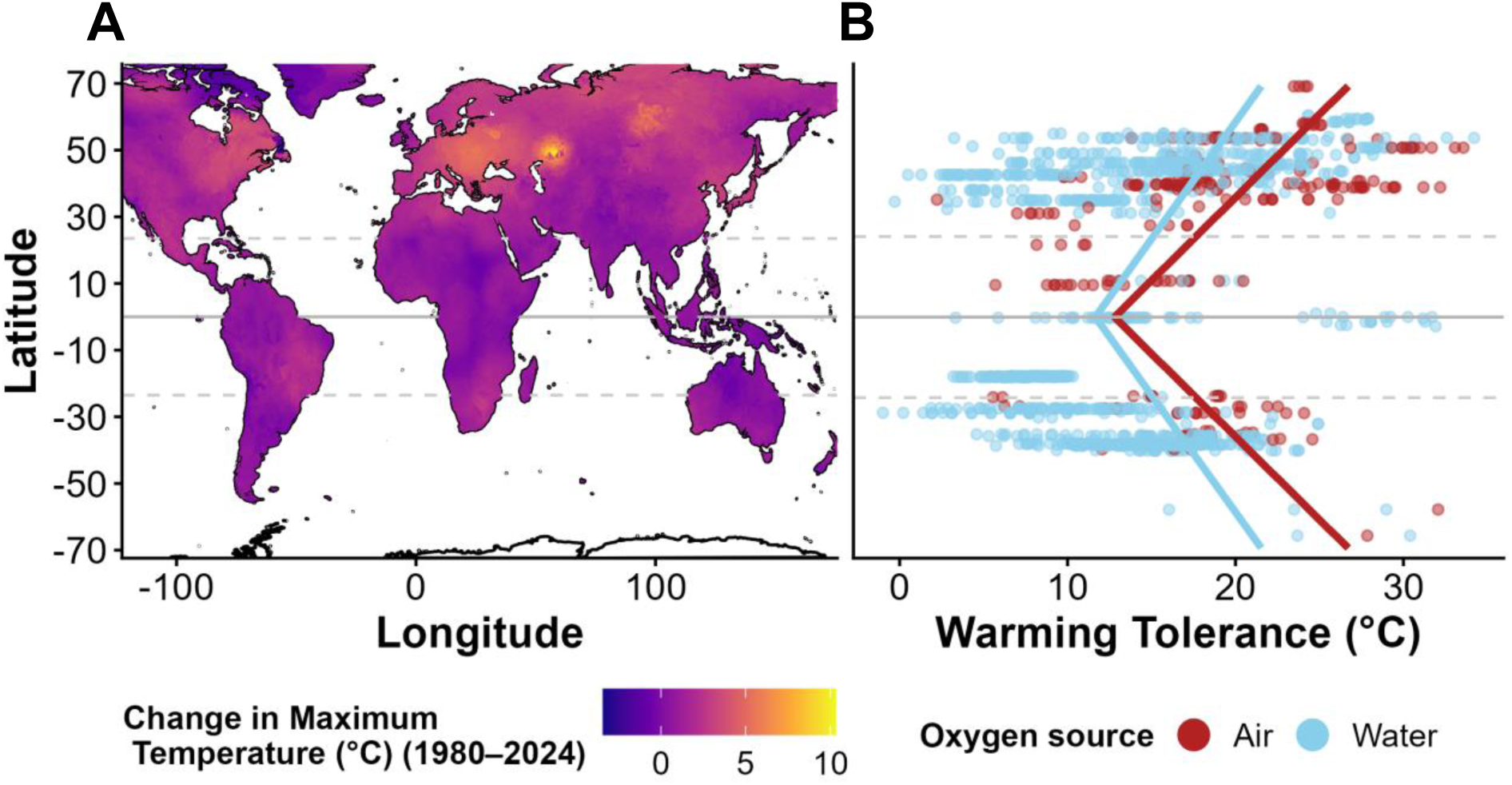
Relationship between species exposure to increasing heat stress and estimated vulnerability. **(A)** Global relationship between latitude and changes in maximum temperatures of aquatic environments from 1980 to 2024. **(B)** PGLMM-estimated slopes of warming tolerance across latitude for species relying on dissolved oxygen (blue) and atmospheric oxygen (red). Warming tolerance was calculated as the difference between upper thermal limits and maximum habitat temperatures.

**Table 3.** PGLMM estimates of oxygen source (air vs water), absolute latitude, and their interactive effects on warming tolerance (°C; upper thermal limit - maximum habitat temperature)

| Term |  | Estimate | S.E. | Z | p-value |
| --- | --- | --- | --- | --- | --- |
| Oxygen source | **Water (intercept)** | **11.66** | **5.494** | **2.122** | **0.034** |
|  | Air | 1.14 | 1.280 | 0.888 | 0.374 |
| **Latitude (Water)** |  | **0.15** | **0.018** | **8.491** | **<0.001** |
| Latitude * Oxygen source (Air) |  | 0.06 | 0.033 | 1.844 | 0.065 |

## Discussion

We show that aquatic insect UTLs are primarily shaped by maximum habitat temperatures and factors related to environmental oxygen availability and its uptake. Variables linked to oxygen availability, including oxygen partial pressure (pO_2_), habitat types, i.e., lotic (high oxygen) or lentic (low oxygen), and insect breathing mode were all strong predictors of UTLs. Together, these results support both the Climate Extremes Hypothesis and the Oxygen- and Capacity-Limited Thermal Tolerance hypothesis. From a trait perspective, functional feeding group explained additional variation in UTLs, while body size did not. When linking UTLs to maximum habitat temperatures to assess warming tolerance, aquatic insects relying on dissolved oxygen experience habitat temperatures closer to their UTLs than taxa with access to atmospheric air, suggesting greater vulnerability for these species or life stages under warming. Additionally, acclimation responses indicate that short-term heat shocks, such as those during mild heat waves, may temporarily increase heat tolerance but provide limited buffering against chronic warming. We further discuss the mechanisms, implications, and caveats underlying these patterns below.

### Support for the Climate Extremes and Oxygen-Limitation Hypotheses

Thermal extremes have long been predicted to drive thermal tolerance rather than average conditions (Bozinovic, 2011; Pither, 2003). Latitude and elevation are commonly used as proxies for climatic gradients, and studies exploring large-scale patterns of UTLs have shown that taxa from temperate latitudes and low elevations tend to have higher UTLs than those from tropical and high-elevation environments (Dewenter et al. 2024; Garcia-Robledo et al. 2020; Shah, Gill, et al. 2017; Dewenter et al. 2025). In our analysis, UTLs increased with absolute latitude, but were most strongly related to maximum water temperature, underscoring the benefit of using actual habitat temperatures over geographic proxies (Verberk et al., 2026). Even infrequent exposure to extreme temperatures may therefore impose strong selection for elevated heat tolerance, as documented more broadly across ectotherms (Sunday et al., 2019).

Warming increases metabolic demand for oxygen, and traits linked to oxygen acquisition were often the strongest predictors of UTLs. Breathing mode—the mechanism by which insects acquire oxygen—was strongly correlated with UTL variation. Insects that are obligate dissolved-oxygen breathers (i.e., gill- or cuticle-breathing insects) displayed the lowest UTLs. The differences in UTLs between life stages likely reflect corresponding shifts in breathing mode, as many hemimetabolous insects are cuticular or gill-breathing juveniles but air-breathing adults (**Appendix 6, Fig. S4;** Buchwalter et al. 2019). Consequently, juvenile life stages dependent on dissolved oxygen are likely to face heightened vulnerability, even in species whose adults tolerate high temperatures (e.g., most species within Odonata, Ephemeroptera, Plecoptera, and Trichoptera).

Compared to air, breathing underwater is challenging. Rates of oxygen diffusion are much lower in water, and these difficulties are compounded by the effects of a higher density and viscosity of water (Verberk et al. 2011; Verberk & Atkinson, 2013). At high elevations, low atmospheric pressure reduces availability of oxygen, which we found to be weakly associated with lower UTLs. Although elevation values were generally similar between collection and test locations, high-elevation populations were, in some cases, tested at lower elevations, which may reduce apparent oxygen limitation during assays and contribute to the weak effect observed for pO_2_. Oxygen limitation may therefore contribute to variation in UTLs among aquatic insects that rely exclusively on dissolved oxygen (Verberk et al., 2016; Verberk & Bilton, 2013).

The combined effects of temperature and oxygen availability on UTLs lead to the prediction that species from lotic (running-water) habitats should exhibit lower heat tolerance and greater sensitivity to hypoxia than species from lentic (standing-water) habitats, as these environments often differ in their thermal and oxygen regimes (Salem, 2021). These patterns of heat tolerance and oxygen sensitivity have been reported for lotic and lentic species of peracarid crustaceans (Verberk et al., 2018) and stoneflies (Frakes et al., 2021). Similarly, we found lentic insects exhibited UTLs ∼ 8 °C higher than their lotic counterparts in independent comparisons of UTLs by habitat type (**Appendix 8; Table S9**). However, once phylogeny, environmental conditions, and organismal traits were incorporated, differences in UTLs became minimal, suggesting that apparent differences between lentic and lotic systems largely reflect variation in environmental temperature, oxygen availability, and taxonomic differences in insect assemblages (Heino et al., 2009; Merritt & Cummins, 1995; Ribera, 2008). In particular, lentic and lotic environments harbor distinct insect communities dominated by different orders, which themselves differ in respiratory strategies and thermal tolerances. In our dataset, around 90% of lotic species relied primarily on dissolved oxygen via cuticular or gill respiration, compared to 64% of lentic species. Differences in temperature and oxygen regimes between lentic and lotic habitats have likely contributed to this taxonomic and trait differentiation (see **Appendix 13** for comparisons of maximum habitat temperatures and pO_2_ across habitat types). Future research should examine how joint warming and deoxygenation may restructure lentic and lotic communities by selectively favoring more tolerant taxa, potentially amplifying existing habitat-associated differences in community composition.

### Differences in UTLS based on functional feeding groups

Although the mechanisms underlying variation in UTLs among functional feeding groups remain uncertain, they likely reflect both trait-mediated physiological differences among feeding guilds and variation in thermal exposure across preferred microhabitats. Scrapers and shredders, including many Ephemeroptera, Trichoptera, and Plecoptera, exhibited some of the lowest UTLs. These groups are typically associated with cool, well-oxygenated environments and rely on biofilm grazing or shredding leaf litter (Jourdan et al., 2018; Pyne & Poff, 2017; Szałkiewicz et al., 2022; Vannote et al., 1980), suggesting that specialization of consumers to food resources available in cool microhabitats may constrain their tolerance and persistence under future warming. Corroborating this finding, Pyne and Poff (2017) identified shredders and biofilm grazers to be among the most vulnerable functional groups under future warming, with declines in species richness reaching 40% in some scenarios. By contrast, filterers and piercer-herbivores exhibited comparatively higher UTLs (Tomczyk et al., 2022). Many filterers, such as mosquito larvae, disproportionately occur in warm and often ephemeral lentic habitats, or downstream in lotic systems, with higher mean temperatures (Vannote et al., 1980). Similarly, piercer-herbivores may occupy warmer, littoral zones of lentic environments while feeding on macrophytes (Merritt et al., 2019). Nevertheless, substantial overlap in UTLs among FFGs indicates that feeding mode alone does not fully drive variation in heat tolerance. Additionally, this overlap could reflect both coarse (e.g., family level) FFG classifications (Kefford et al., 2020; Merritt & Cummins, 1995), and overlap between feeding guilds, breathing mode, and phylogeny among taxa in our analysis, validating our approach of combining these factors into a single model.

### Effects of acclimation temperature and duration on UTLs

Many species in our dataset were acclimated for hours to a few days; durations that typically elicit passive and immediate responses to warming (i.e., Q_10_ effects) rather than true acclimation involving physiological adjustments, which occurs over longer timescales (Havird et al., 2020). In line with these expectations, brief exposure to elevated temperatures (up to ∼7 days above species-specific mean acclimation temperatures) resulted in higher UTLs, whereas longer periods (e.g., >3 weeks) consistently resulted in lower UTLs, suggesting that insects experience cumulative thermal damage under prolonged exposure (Loeschcke & Hoffmann, 2002; Lutterschmidt & Hutchison, 1997; Ørsted et al., 2022; Terblanche et al., 2007). Because thermal injury accumulates over time, species that appear tolerant in short-term assays may still experience substantial sublethal stress or mortality under prolonged environmental exposure not captured by acute heat tolerance estimates like UTLs (Jørgensen et al., 2021; Ørsted et al., 2022; Rezende et al., 2014). Future work should therefore evaluate how acclimation effects interact with heat intensity and exposure duration to shape thermal limits under realistic climate scenarios (Verberk et al., 2023).

### How vulnerable are aquatic insects to climate change?

Across taxa, warming tolerance was lowest in tropical regions, where species already live closest to their thermal limits. This finding is consistent with studies predicting greater vulnerability in tropical regions (Deutsch et al., 2008; Pinsky et al., 2019; Shah, Gill, et al., 2017; Sheldon et al., 2011). However, our results further highlight the importance of breathing mode and oxygen limitation in shaping vulnerability across freshwater taxa. Dissolved-oxygen breathers tend to live closer to their UTLs, and their warming tolerance increases more slowly with latitude than taxa with access to atmospheric oxygen, suggesting that reliance on dissolved oxygen may constrain thermal safety margins toward mid- and high latitudes. Although warming tolerance is lowest in tropical regions, rapid warming and increasing heat-wave intensity at mid- and high latitudes may in fact, significantly increase vulnerability of temperate freshwater assemblages (Bachmann, 2025; Piccolroaz et al., 2020; Pinsky et al., 2019). Indeed, our findings identify reliance on dissolved oxygen, and thus, oxygen availability, as a key constraint on tolerance to heat stress, placing most aquatic insects at heightened risk under climate warming.

Because of the influence of oxygen availability on UTLs, vulnerability is likely amplified in aquatic life stages. Juvenile aquatic insects typically exhibit lower UTLs and rely more strongly on dissolved oxygen than terrestrial adults (e.g., Odonata, Ephemeroptera, Plecoptera, and Trichoptera), and many aquatic insects spend the majority of their life cycle as juveniles. As a result, even taxa with relatively heat-tolerant adults may remain at risk when heat waves occur during the more sensitive juvenile stages. Such constraints may help explain projections of higher extinction risk for aquatic insects (33%) compared to terrestrial insects (28%) under future warming scenarios (Cardoso et al., 2020; Harvey et al., 2023; Sánchez-Bayo & Wyckhuys, 2019).

Our rapidly warming global climate is sparking more frequent, intense, and prolonged heat waves (Meehl & Tebaldi, 2004). Although some degree of plasticity in thermal limits may offer initial buffering against short term warming, the capacity for aquatic insects to adjust UTLs appears limited, and empirical evidence remains mixed across taxa (Weaving et al., 2022). For example, temperate mayflies from different elevations in the Rocky Mountains exhibited plasticity in CT_MAX_ following short-term warming, whereas co-occurring stoneflies did not (Birrell et al., 2023; Shah, Funk, et al., 2017). More broadly, there is limited evidence that physiological plasticity or evolutionary adaptation can keep pace with ongoing rates of climate warming (Gunderson & Stillman, 2015; Hoffmann & Sgrò, 2011). Together, these patterns indicate that aquatic insect vulnerability will be driven not only by acute thermal extremes, but also by sustained exposure to warming and deoxygenation, which can impair growth, emergence success, and recruitment well before UTLs are reached (Shah et al., 2023). These findings highlight the need to integrate thermal limits with sublethal performance metrics and recovery dynamics when assessing patterns of vulnerability.

### Conclusion and future directions

Our synthesis demonstrates that multiple, non-mutually exclusive factors shape variation in UTLs across aquatic insects. Among them, maximum habitat temperature and oxygen availability are powerful predictors of UTLs, revealing that many aquatic insects already operate close to their physiological limits. While warming tolerance increased with latitude, temperatures are rising most rapidly at mid- and high-latitudes (IPCC, 2021) indicating that the wider thermal safety margins of temperate species relying on dissolved oxygen may contract as rapid, unmitigated warming continues. Moving forward, integrating high-resolution physiological data with local climatic conditions, organismal traits, and phylogenetic context will be essential not only for identifying vulnerable species and communities but also for predicting how aquatic ecosystem structure and function may shift under climate warming.

## Acknowledgements

The late BSD contributed data for seasonal and acclimatory effects on UTLs using Australian species of Ephemeroptera, Plecoptera, and Trichoptera. We gratefully acknowledge the authors of previously published upper thermal limit (UTL) datasets that contributed to this synthesis, including Bayat et al. (2025), Bennett et al. (2021), Leiva et al. (2019), Chown et al. (2015), and Weaving et al. (2022).

