## Supporting Information for "Global synthesis of aquatic insect heat tolerance reveals oxygen availability as a key driver of climate vulnerability"

### **Appendix 1**: PRISMA literature review

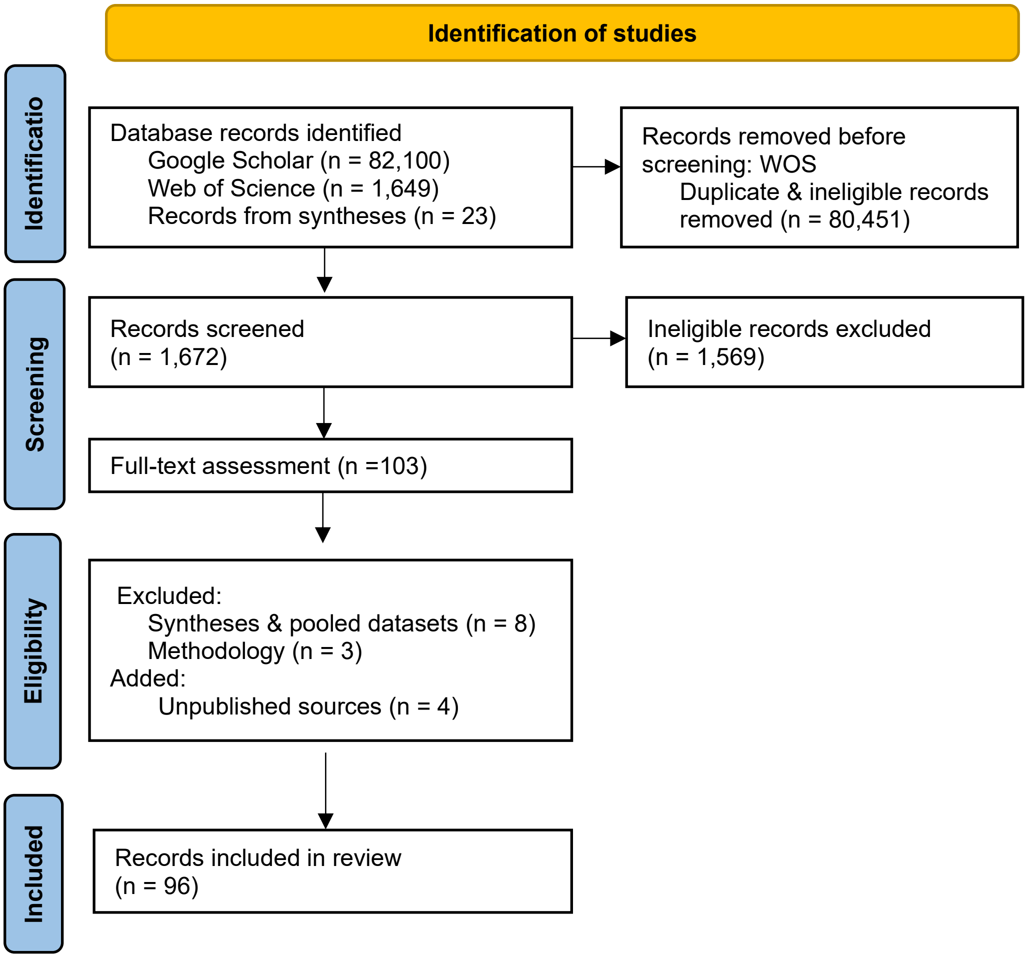

**Figure S1**. Preferred Reporting Items for Systematic reviews and Meta-Analyses (PRISMA). Overview of data sources identified by search terms (described in methods) and exclusion process in manual review (Page et al. 2021).

### **Appendix 2:** Time-standardized correction for thermal limits

Results from two-way ANOVAs comparing upper thermal limits across dynamic ramping and static assay types, before correction using raw thermal limits and corrected values standardized to a 100-minute assay duration.

**Table S1**: Effects of thermal assay type on upper thermal limits before and after standardization

| Response variable | Effect | df | Sum of squares | Mean square | F | p |
|---|---|---|---|---|---|---|
| Thermal limits | Assay type | 1 | 13,618 | 13,618 | 350.2 | < 0.001 |
| | Residuals | 1,344 | 52,268 | 38.9 | — | — |
| Corrected thermal limits (UTL) | Assay type | 1 | 345.3 | 345.3 | 9.30 | 0.002 |
| | Residuals | 1,344 | 49,913 | 37.1 | — | — |

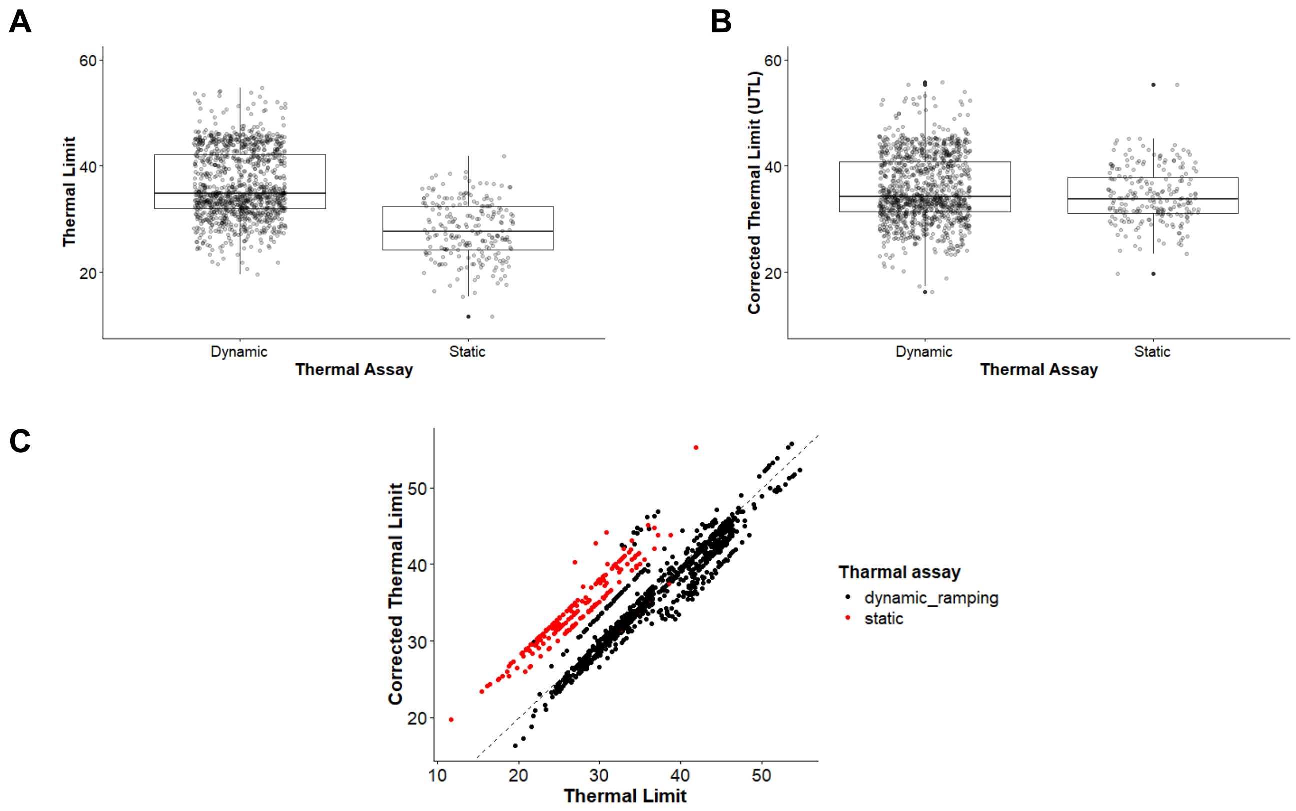

**Figure S2**. Distribution of thermal limits measured using dynamic ramping and static assays **(A)** before and **(B)** after 100 minute time standardized correction (Corrected Thermal Limits). **(C)** Relationship between thermal limits to corrected limits (UTLs) which were used throughout analyses as the response variable.

#### **Appendix 3:** Distributions of assay types across taxa and oxygen sources

Assay type was not randomly distributed across taxa or oxygen source. Static assays were used almost exclusively on taxa that rely on dissolved oxygen in water, whereas dynamic ramping assays included a higher proportion of air-breathing taxa ($\chi^2 = 49.7$, df = 1, $p < 0.001$; Table S2).

Similarly, assay type differed strongly among insect orders ($\chi^2 = 130.9$, df = 7, $p < 0.001$), with Ephemeroptera disproportionately represented in static assays and Diptera, Coleoptera, and Odonata more frequently measured using dynamic ramping methods (Table S3)

**Table S2**. Proportion of thermal limit data from dynamic or static assays across by primary oxygen source.

| Thermal assay type | Dynamic ramping | Static |
|---|---|---|
| Air-breathing | 0.226 | 0.018 |
| Water-breathing | 0.774 | 0.982 |

**Table S3.** Proportion of thermal limit data from dynamic or static assays across insect orders.

| Order | Dynamic ramping | Static |
|---|---|---|
| Ephemeroptera | 0.198 | 0.489 |
| Trichoptera | 0.220 | 0.169 |
| Plecoptera | 0.164 | 0.142 |
| Diptera | 0.067 | 0.119 |
| Megaloptera | 0.003 | 0.018 |
| Hemiptera | 0.028 | 0.000 |
| Coleoptera | 0.132 | 0.032 |
| Odonata | 0.188 | 0.032 |

### **Appendix 4:** Upper thermal limits by insect Order

**Generalized linear model (GLM)**

A Gaussian generalized linear model was used to test for differences in upper thermal limits (UTLs) between insect Orders.

**Model structure:**
UTL ~ Order

**Sample size:** 1,346 observations

**Table S4.** Estimated marginal means, standard errors (SE) and 95% confidence intervals (CL) for upper thermal limits of each order

| Order | UTL (°C) | SE | Lower CL | Upper CL |
|---|---|---|---|---|
| Ephemeroptera | 31.6 | 0.226 | 31.1 | 32.0 |
| Trichoptera | 32.3 | 0.243 | 31.8 | 32.7 |
| Plecoptera | 32.3 | 0.279 | 31.8 | 32.9 |
| Diptera | 38.0 | 0.408 | 37.2 | 38.8 |
| Megaloptera | 38.1 | 1.549 | 35.0 | 41.1 |
| Hemiptera | 41.8 | 0.724 | 40.3 | 43.2 |
| Coleoptera | 42.0 | 0.328 | 41.4 | 42.6 |
| Odonata | 41.9 | 0.277 | 41.4 | 42.5 |

### **Appendix 5:** Breathing mode & functional feeding group results

**Table S5.** PGLMM predicted mean UTL estimates, empirical standard errors and 95% confidence intervals for each biotic trait term**.** Results correspond to Fig. 3C & D in manuscript.

| Bioitc trait | Group | N | Estimate | SE | Lower CL | Upper CL |
|---|---|---|---|---|---|---|
| Breathing mode | Gills | 988 | 35.61 | 0.011 | 35.59 | 35.63 |
| | Cuticle | 66 | 36.05 | 0.048 | 35.96 | 36.15 |
| | Plastron | 33 | 39.70 | 0.066 | 39.57 | 39.83 |
| | Gills/Spiracles | 36 | 42.78 | 0.033 | 42.71 | 42.84 |
| | Spiracles | 165 | 43.41 | 0.026 | 43.36 | 43.46 |
| | Siphon | 58 | 43.33 | 0.027 | 43.27 | 43.38 |
| Functional feeding group | Scraper/Shredder | 435 | 35.68 | 0.040 | 35.60 | 35.76 |
| | Gatherer | 169 | 37.00 | 0.062 | 36.88 | 37.13 |
| | Predator | 519 | 37.35 | 0.039 | 37.28 | 37.43 |
| | Filterer | 172 | 40.28 | 0.070 | 40.14 | 40.42 |
| | Piercer–Herbivore | 18 | 41.22 | 0.182 | 40.83 | 41.60 |

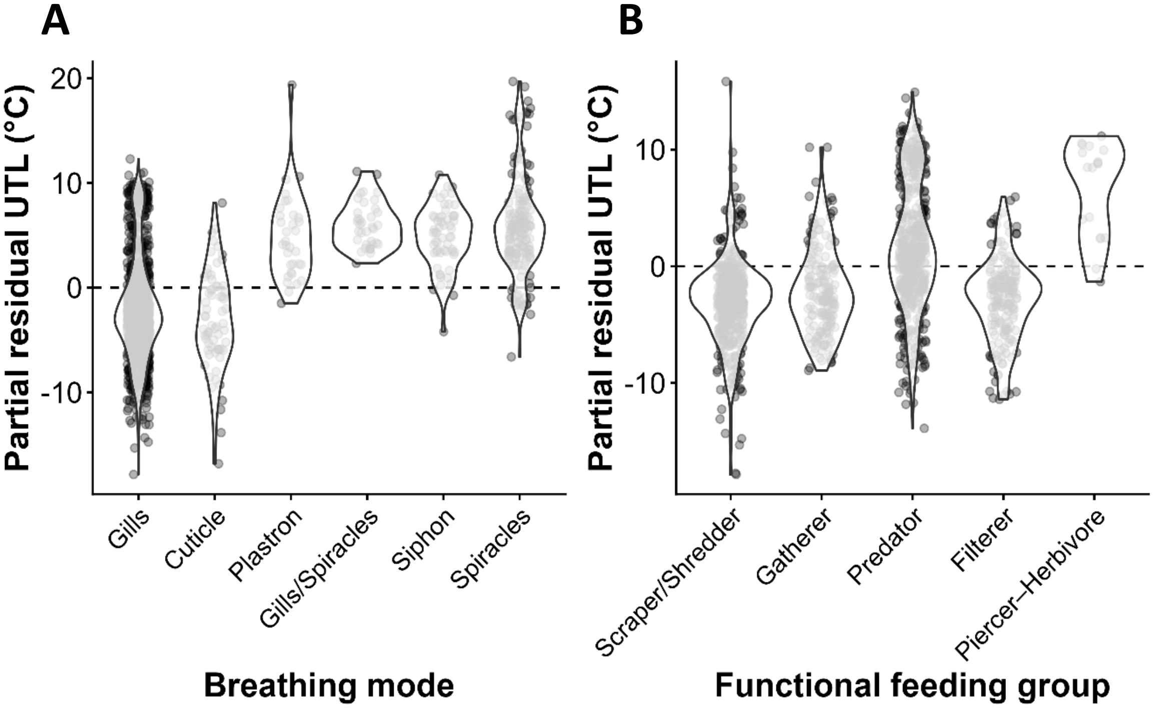

**Figure S3**. Partial residuals from phylogenetic generalized linear mixed models (PGLMMs) illustrating trait effects on upper thermal limits (UTL) after accounting for environmental covariates and the non-focal trait (FFG or breathing mode). Panels show partial residual UTL values for **(A)** breathing modes and **(B)** functional feeding groups. Partial residuals were calculated as observed UTL minus fitted values from models excluding the focal trait, while retaining maximum habitat temperature and oxygen availability ($pO_2$) as covariates. The dashed horizontal line at zero indicates the expected UTL given environmental conditions and the non-focal trait; deviations from zero represent higher or lower UTLs than expected after accounting for external variables. Violin plots show the distribution of partial residuals within each trait group, with points representing individual observations.

### **Appendix 6**: Life stage effects on upper thermal limits

A Gaussian linear model was used to test differences in upper thermal limits (UTLs) between life stages.

**Model structure:**
UTL ~ life stage (adult vs juvenile)

**Sample size:** 1,346 observations

**Table S6.** LM estimates for mean UTLs by life stage

| Predictor | Estimate | SE | t | p |
|---|---|---|---|---|
| Intercept (juvenile) | 34.00 | 0.155 | 219.63 | < 0.001 |
| Life stage (adult) | 8.65 | 0.373 | 23.21 | < 0.001 |

Residual deviance = 35877 (df = 1344), AIC = 8245

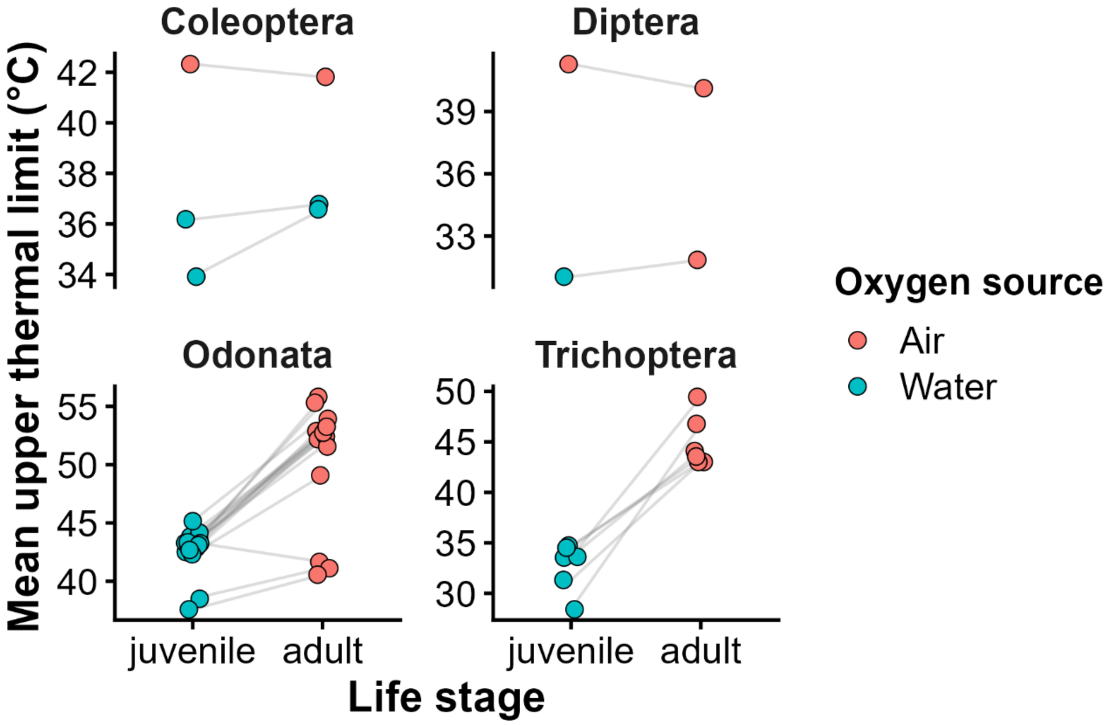

**Figure S4.** Species-level paired comparisons of upper thermal limits by life stage. Each point represents the mean UTL of a life stage for a given species, and grey lines connect juvenile-adult pairs. Panels correspond to insect order and colors to primary oxygen source (air=red & water=blue).

### **Appendix 7**: Body size effects on upper thermal limits

A Gaussian linear model was used to examine the relationship between upper thermal limits (UTLs) and body size ($\log_{10}$-transformed mass), including mass measurement type (wet vs dry) and their interaction.

**Model structure:**
UTL ~ $\log_{10}$(mass) × mass type

**Sample size:** 197 observations

**Table S7.** LM estimates of mean UTLs by mass, mass type, and their interaction

| Predictor | Estimate | SE | t | p |
|---|---|---|---|---|
| Intercept | 33.63 | 0.804 | 41.82 | < 0.001 |
| $\log_{10}$(dry mass) | 2.21 | 0.335 | 6.58 | < 0.001 |
| Mass type (wet mass) | 1.47 | 1.199 | 1.22 | 0.223 |
| $\log_{10}$(mass) × Mass type (wet) | 1.70 | 0.784 | 2.17 | 0.031 |

Residual deviance = 4312.3 (df = 193), AIC = 1177

---

**Linear mixed-effects model (LMER): species identity as a random effect**

To account for non-independence among species with body mass, we fit a linear mixed-effects model with species identity included as a random intercept.

**Sample size:** 197 observations
**Number of species:** 90

**Table S8.** LMM estimates of mean UTLs by mass, measurements type (dry=reference), and their interaction**.** Species identity was included as a random intercept

| Predictor | Estimate | SE | t | p |
|---|---|---|---|---|
| Intercept | 34.40 | 0.913 | 37.66 | <0.001 |
| $\log_{10}$ (dry mass) | 1.48 | 0.359 | 4.14 | <0.001 |
| Mass type (wet) | 0.63 | 1.153 | 0.55 | 0.586 |
| $\log_{10}$(mass) × Mass type (wet) | −0.63 | 0.567 | −1.11 | 0.272 |

#### **Appendix 8**: Habitat type effects on upper thermal limits

A Gaussian linear model was used to examine differences in upper thermal limits (UTLs) between non-terrestrial habitat types, comparing lotic and lentic systems, and excluding data from adult life stages of species with terrestrial adults.

**Model structure:**
UTL ~ habitat type (lentic vs lotic)

**Sample size:** 1,282 observations

**Table S9.** LM estimates of mean UTLs by lotic (reference) and lentic habitat types

| Predictor | Estimate | SE | t | p |
|---|---|---|---|---|
| Intercept (lotic) | 33.50 | 0.153 | 218.59 | < 0.001 |
| Habitat type (lentic) | 8.08 | 0.346 | 23.36 | < 0.001 |

Residual deviance = 30966 (df = 1280), AIC = 7727

### **Appendix 9**: Acclimation effects on upper thermal limits

A gaussian linear model was used to visualize acclimation effects without phylogenetic structure, using standardized acclimation temperature, acclimation duration, and their interaction.

**Model structure:**
UTL ~ standardized acclimation temperature * acclimation duration

**Sample size:** 1,236 observations

**Table S10**. LM results for effects of standardized acclimation temperature (Δ°C; acclimation temperature - species-specific mean acclimation temperature), acclimation duration ($log_{10}$ hours), and their interaction on upper thermal limits.

| Predictor | Estimate | SE | t | p |
|---|---|---|---|---|
| Intercept | 26.78 | 0.613 | 43.72 | <0.001 |
| Standardized acclimation temperature | 0.70 | 0.163 | 4.28 | <0.001 |
| Acclimation duration ($log_{10}$ hours) | 4.87 | 0.343 | 14.17 | <0.001 |
| Standardized acclimation temperature × Acclimation duration ($log_{10}$ hours) | −0.34 | 0.093 | −3.70 | <0.001 |

Residual deviance = 6426 (df = 1233), AIC = 5559

### **Appendix 10**: Warming Tolerance

Warming tolerance was quantified as the difference between upper thermal limits and maximum environmental temperature. A gaussian linear model was fit for warming tolerance as a function of absolute latitude and primary oxygen source (i.e., water or air).

**Model structure:**
warming tolerance ~ absolute latitude * oxygen source

**Sample size:** 1,346 observations

**Table S11**. LM results for warming tolerance across absolute latitude, oxygen source (air=reference), and their interaction.

| Predictor | Estimate | SE | t | p |
|---|---|---|---|---|
| Intercept (Air) | 9.76 | 1.112 | 8.78 | < 0.001 |
| Absolute latitude | 0.26 | 0.029 | 8.88 | < 0.001 |
| Oxygen source (water) | −1.52 | 1.218 | −1.25 | 0.211 |
| Absolute latitude × oxygen source (water) | −0.09 | 0.032 | −2.88 | 0.004 |

Residual deviance = 45240 (df = 1,342), AIC = 8561

### **Appendix 11**: Global map of collection sites

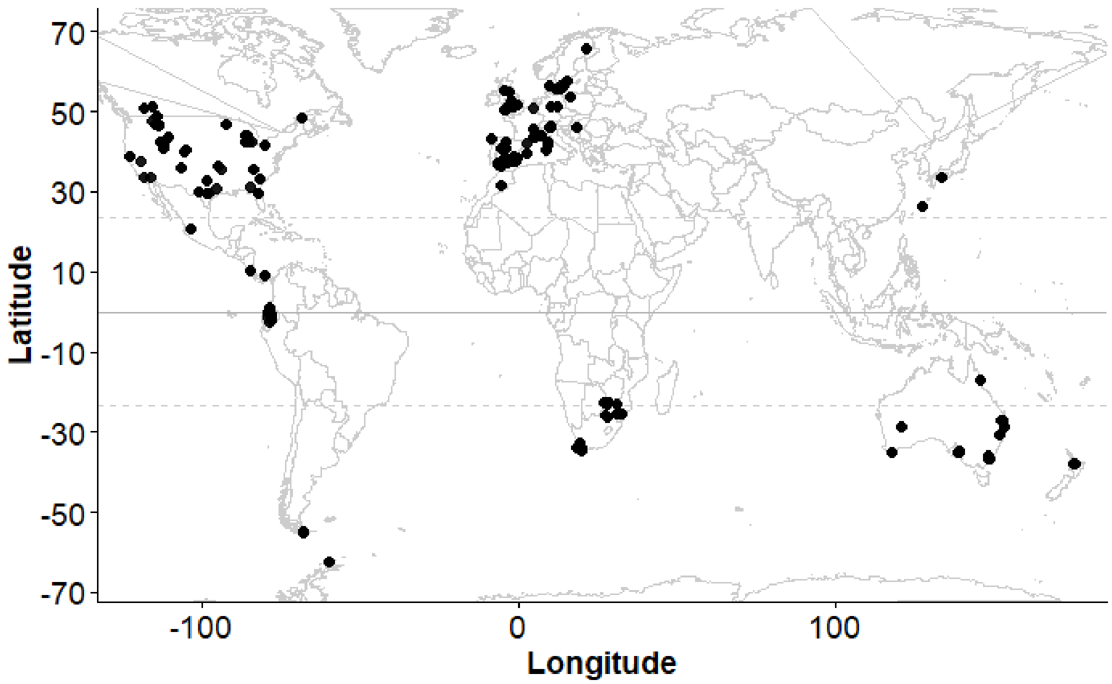

**Figure S5**. Global map of unique coordinates of collection sites for insects in the dataset

### **Appendix 12**: Maximum habitat temperature by oxygen source

A gaussian linear model fit on maximum habitat temperature as a function of primary oxygen source (i.e., water or air) was used to compare estimates of maximum habitat temperatures for terrestrial or aquatic life stages of organisms in our dataset.

**Model structure:**
Maximum habitat temperature ~ oxygen source

**Sample size:** 1,346 observations

**Table S12**. LM results for maximum habitat temperature across oxygen sources (air=reference).

| Predictor | Estimate | SE | t | p |
|---|---|---|---|---|
| Intercept (Air) | 23.35 | 0.289 | 80.76 | < 0.001 |
| Oxygen source (water) | -3.42 | 0.322 | −10.62 | < 0.001 |

Residual deviance = 29109 (df = 1,344), AIC = 7963

### **Appendix 13**: Maximum habitat temperature and $pO_2$ by habitat types

Gaussian linear models were used to examine differences in maximum habitat temperatures and $pO_2$ between non-terrestrial habitat types, comparing lotic and lentic systems.

**Sample size:** 1346 observations

**Model structure:**
Maximum habitat temperature ~ habitat type (lentic vs lotic)

**Table S13.** LM estimates for mean maximum habitat temperature by lotic (reference) and lentic habitat types

| Predictor | Estimate | SE | t | p |
|---|---|---|---|---|
| Intercept (lotic) | 20.11 | 0.143 | 140.99 | < 0.001 |
| Habitat type (lentic) | 0.75 | 0.322 | 2.34 | 0.019 |

Residual deviance = 26811 (df = 1280), AIC = 7542

**Model structure:**
$pO_2$ ~ habitat type (lentic vs lotic)

**Table S14.** LM estimates for mean $pO_2$ by lotic (reference) and lentic habitat types

| Predictor | Estimate | SE | t | p |
|---|---|---|---|---|
| Intercept (lotic) | 19.46 | 0.053 | 368.10 | < 0.001 |
| Habitat type (lentic) | 0.73 | 0.119 | 6.148 | < 0.001 |

Residual deviance = 3684 (df = 1280), AIC = 4997
